# Towards a Survival Risk Prediction Model for Metastatic NSCLC Patients on Durvalumab Using Whole-Lung CT Radiomics

**DOI:** 10.1101/2024.02.01.578458

**Authors:** Kedar Patwardhan, Harish RaviPrakash, Nikos Nikolaou, Ignacio Gonzalez-García, José Domingo Salazar, Paul Metcalfe, Joachim Reischl

## Abstract

**Background:** Existing criteria for predicting patient survival from immunotherapy are primarily centered on the PD-L1 status of patients. We tested the hypothesis that noninvasively captured baseline whole-lung radiomics features from CT images, baseline clinical parameters, combined with advanced machine learning approaches, can help to build models of patient survival that compare favorably with PD-L1 status for predicting ‘less-than-median-survival risk’ in the metastatic NSCLC setting for patients on durvalumab. With a total of 1062 patients, inclusive of model training and validation, this is the largest such study yet.

**Methods:** To ensure a sufficient sample size, we combined data from treatment arms of three metastatic NSCLC studies. About 80% of this data was used for model training, and the remainder was held-out for validation. We first trained two independent models; Model-C trained to predict survival using clinical data; and Model-R trained to predict survival using whole-lung radiomics features. Finally, we created Model-C+R which leveraged both clinical and radiomics features.

**Results:** The classification accuracy (for median survival) of Model-C, Model-R, and Model-C+R was 63%, 55%, and 68% respectively. Sensitivity analysis of survival prediction across different training and validation cohorts showed concordance indices ([95 percentile]) of 0.64 ([0.63, 0.65]), 0.60 ([0.59, 0.60]), and 0.66 ([0.65,0.67]), respectively. We additionally evaluated generalization of these models on a comparable cohort of 144 patients from an independent study, demonstrating classification accuracies of 65%, 62%, and 72% respectively.

**Conclusion:** Machine Learning models combining baseline whole-lung CT radiomic and clinical features may be a useful tool for patient selection in immunotherapy. Further validation through prospective studies is needed.

## 1 Introduction

Lung cancer is the leading cause of cancer death and makes up around 25% of cancer deaths worldwide (1). The 5-year survival for patients with non-small cell lung cancer (NSCLC) is approximately 25% but drops to almost 7% in the case of metastatic NSCLC (mNSCLC). Recent advances have led to development of targeted immune checkpoint inhibitors such as durvalumab, targeting programmed death ligand 1 (PD-L1), which has shown durable clinical benefit to patients with locally advanced NSCLC (2). In this case levels of PD-L1 expression, derived from evaluation of biopsy tissue, have been shown to impact overall survival (OS) following durvalumab treatment (3). Lung biopsies are invasive, often challenging due to anatomical access, do not account for inter/intra lesion heterogeneity, and are unable to provide evaluations for all lesions. Radiological imaging e.g., computed tomography (CT), provide non-invasive characterization of all lesions as well as the surrounding anatomical region. This information is available at baseline as well as longitudinal time-points, whereas longitudinal biopsies are impractical. This has been a key motivation for our exploration of non-invasive imaging based patient stratification approaches in the mNSCLC setting.

### Literature Review

Given that OS is typically a primary endpoint for clinical trials to demonstrate efficacy, we searched the PubMed database to identify relevant prior work in the field of mNSCLC where baseline CT images (radiomics) were used for OS prediction in patients on immunotherapy (IO) treatment (4). The most relevant publications as of December 1, 2022, are highlighted in Table 1^1^.

**Table 1.** Overview of Relevant Prior Work Towards OS Prediction Using Baseline CT Images in Patients on IO, in a mNSCLC Setting. Values not reported are indicated with N/A. N_test_ indicates the number of “held-out” samples used to test a model (these samples are not seen by the model during the training phase).

| Reference | Total no. of Patients | Accuracy (%) in Test Cohort<br>(mutually exclusive training and test sets sampled from same cohort) | AUC in Test Cohort<br>(mutually exclusive training and test sets sampled from same cohort) | Concordance Index (robustness to training/test splits) | Generalization Accuracy (%) (independent cohort) |
| --- | --- | --- | --- | --- | --- |
| Jazieh et al (20) | 133 | N/A ($N_{\text{test}} = 69$) | N/A | 0.69 (N/A) | N/A |
| Gong et al (21) | 224 | 50.79 ($N_{\text{test}} = 63$) | 0.52 (0.39–0.65) | 0.51 (0.43–0.6) | 52.94 ($N_{\text{test}} = 68$) |
| He et al (22) | 236 | 83.3 ($N_{\text{test}} = 48$) | N/A | 0.75 (0.59–0.90) | N/A |
| Liu et al (23) | 46 | N/A ($N_{\text{test}} = 1$) | 0.64 (0.48–0.79) | 0.86 (0.74–0.97) | N/A |
| Trebeschi et al (24) | 123 | N/A ($N_{\text{test}} = 42$) | 0.76 (N/A) | N/A | N/A |
| Work Presented* | 917 | 66.8 ($N_{\text{test}} = 199$) | 0.71 (0.65–0.78) | 0.66 (0.65–0.67) | 73.5 ($N_{\text{test}} = 144$) |

Values not reported are indicated with N/A. $N_{\text{test}}$ indicates the number of “held-out” samples used to test a model (these samples are not seen by the model during the training phase).
\*Results reported are for Model-C+R. Evaluation metrics can be found in Supplementary Table 5.
AUC, area under the curve; CT, computed tomography; IO, immunotherapy; OS, overall survival; mNSCLC, metastatic non-small cell lung cancer; NA, not applicable.

While such prior work has demonstrated the benefit of CT radiomics and clinical models, unlike our work, all the models reported in literature have been trained on relatively small datasets of <200 patients, which can limit the variability encountered and thereby model generalization in unseen datasets. More importantly, all prior work requires accurate delineation of the target tumor. This typically requires a slice-by-slice annotation by an experienced radiologist which is a time-intensive task. Such a primary-lesion (and/or tumor macro-environment) focused approach does not take into consideration the contribution of other lesions, anatomical structures (e.g., lymph nodes), as well as the characteristics of healthy lung tissue that may be relevant to the OS of the patients (e.g., overall health status, tolerance to IO, etc).

### Summary of Contributions

In this study we aim to establish the value of non-invasive CT radiomic features in patient selection for durvalumab monotherapy or in combination in an mNSCLC setting. This is the largest study so far looking at baseline CT radiomic features for IO survival prediction in mNSCLC (N > 1000 including training, testing, and independent cohort evaluation). Apart from the sample size, since our data covers multiple sites and clinical trials, there is reasonable diversity in the imaging data, as shown in Table 2, to ensure that models developed are robust to such variations. Additionally, compared to existing literature, we do not use any lesion annotation for extraction of radiomic features while demonstrating favorable performance compared to prior art for survival-based patient risk prediction. The following sections will describe in detail the data, methods, and results, followed by a brief discussion towards next steps in our research.

**Table 2.**
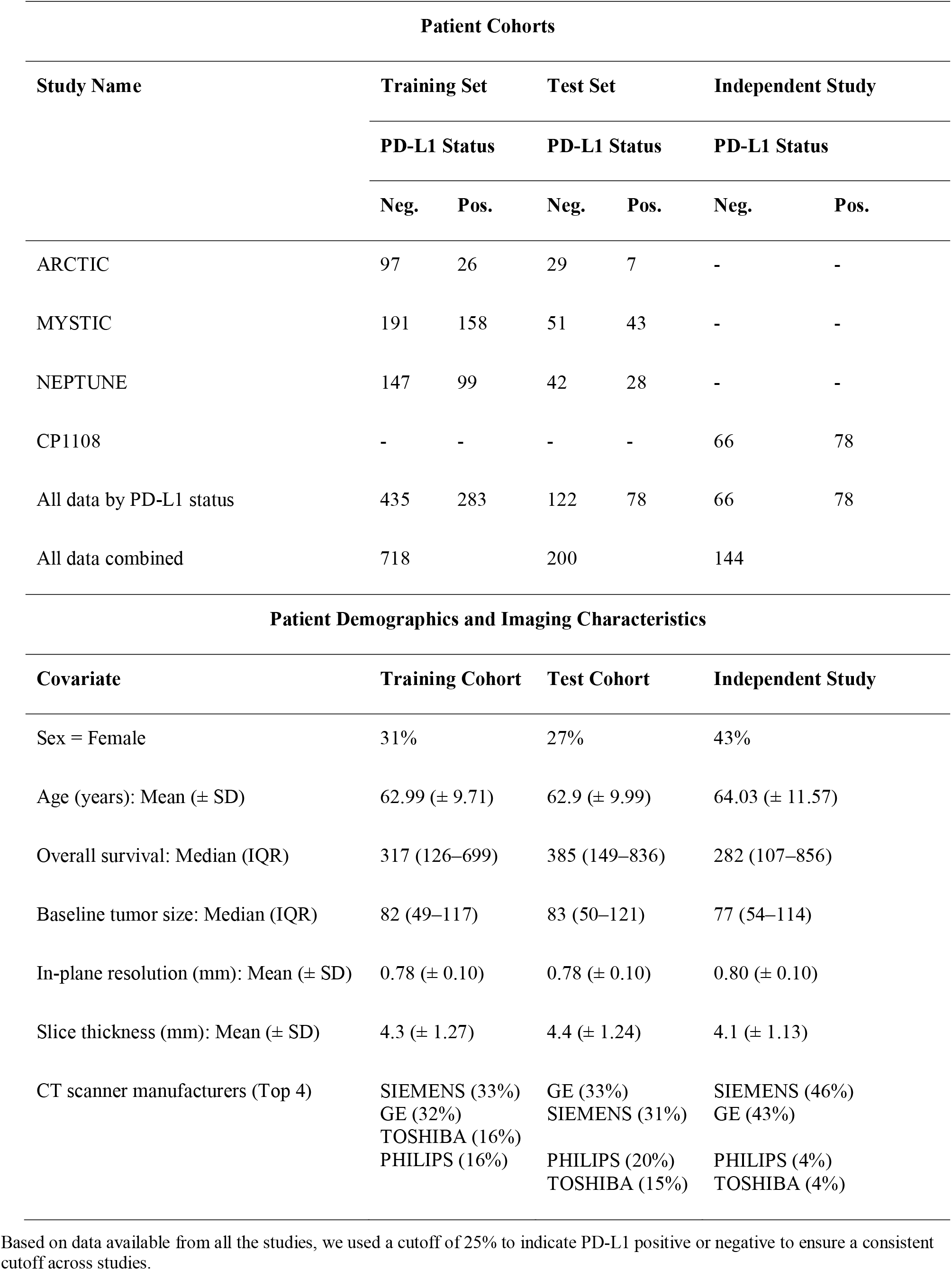
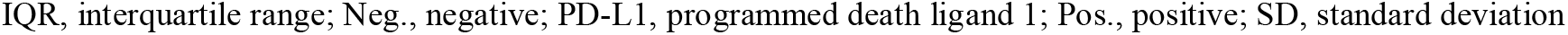
Patient Cohorts and Demographics. Based on data available from all the studies, we used a cutoff of 25% to indicate PD-L1 positive or negative to ensure a consistent cutoff across studies.

## 2 Material and Methods

### 2.1 Data Preparation

We performed a retrospective analysis using data from three phase III clinical studies (ARCTIC [NCT02352948] (5, 6), MYSTIC [NCT02125461] (3, 7), and NEPTUNE [NCT04571658] (8, 9)) investigating durvalumab alone or in combination in patients with a diagnosis of mNSCLC. The analysis presented in the manuscript uses data from patients that provided informed consent including for secondary reuse. All studies were conducted in compliance with the Declaration of Helsinki and the United States (US) Food and Drug Administration (FDA) Guidelines for Good Clinical Practice.

The dataset of 1320 patients were assessed for eligibility. We manually reviewed the images as a quality check for image resolution and motion artifacts and excluded patients with in-plane resolution of over 1 mm or slice thickness (out-of-plane resolution) over 10 mm (n = 107). Finally, patients with missing clinical parameters were also excluded (n = 295) resulting in a total of 918 patients as shown in Supplementary Figure 1. A complete list of the included clinical parameters can be found in the supplementary material (see Supplementary Table 3).

Prior to splitting the analysis data into training and test cohorts, Kaplan-Meier (KM) (10) survival curve of the treatment arms of the three studies was generated (see Supplementary Figure 2a) and no significant difference was observed in the survival times when analyzed as a combined cohort (though it must be noted that there are patient subpopulations where there are differences in survival) (3, 11). Thus, data from the three clinical trials were combined to create a large analysis cohort (referred to as the primary study cohort) that comprised of 918 patients. PD-L1 status for each patient was determined at 25% cut-off^2^. Patients with over 25% PD-L1 expression were labeled as PD-L1-positive and PD-L1-negative otherwise. KM curve for PD-L1 status showed a significant difference between PD-L1-positive and negative patients (Supplementary Figure 2b). The patient cohorts were thus generated via stratifying based on PD-L1 and study, and were split 80:20 into training and test datasets.

In order to use a completely blinded cohort (previously unseen by the model) for validation, a matching cohort (with respect to drug regimen and indication) was selected from a drug-escalation study CP-1108 (NCT01693562) (12) (n = 224). Patients with disease stage III and below at the time of entry to the clinical trial were excluded from the analysis. Patients were further excluded based on the criteria used in the primary cohort. The final independent validation cohort consisted of 144 patients. Number of patients in each study and the PD-L1 distribution is shown in Figure 1a.

**Figure 1.**
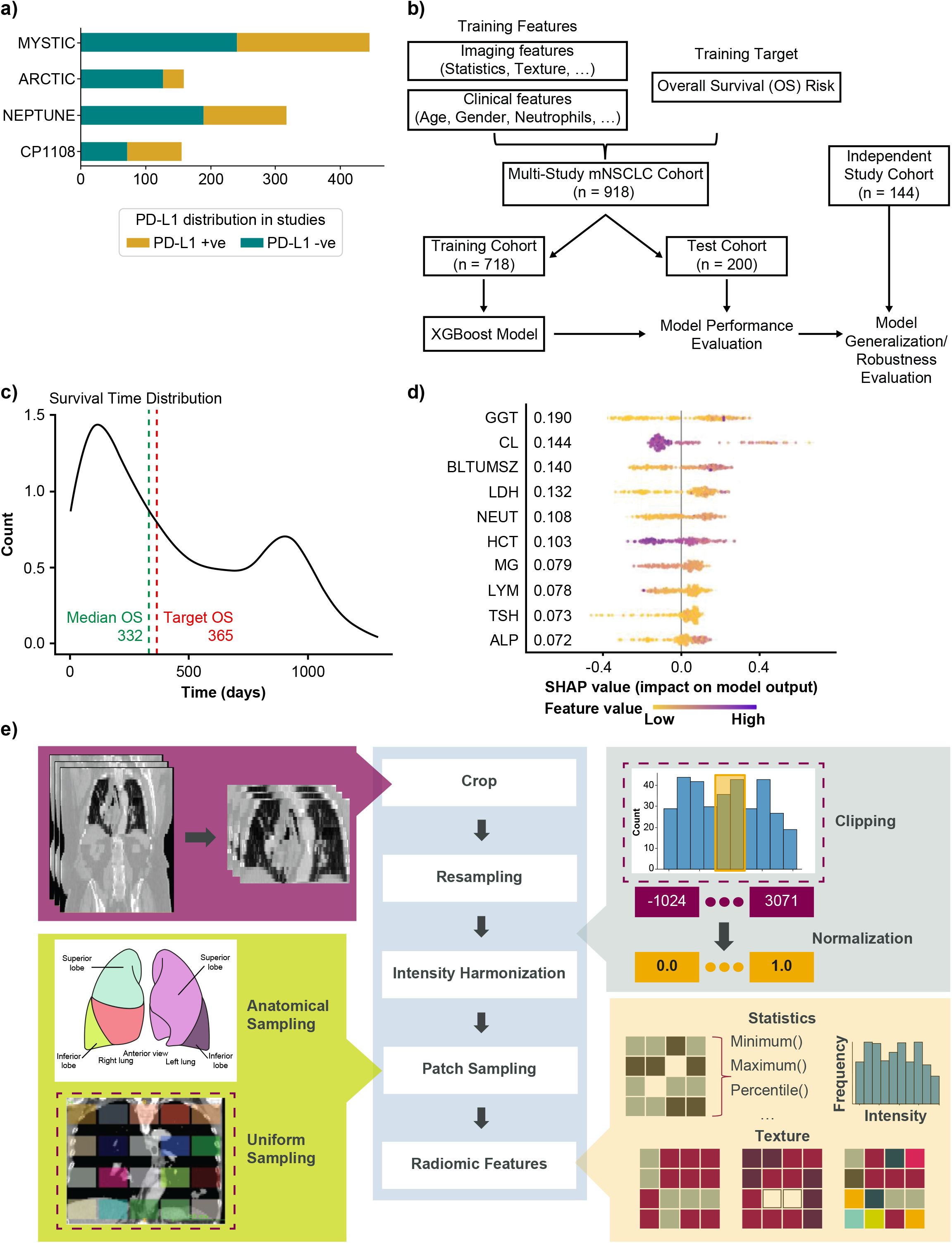
Overview of the modeling for the generation of whole-lung radiomics for survival risk prediction. **a)** Bar chart showing the number of patients in each of the clinical trials. PD-L1 status was determined at 25% cutoff. **b)** Overview of the training and testing procedure using XGBoost model. The MYSTIC, ARCTIC, and NEPTUNE datasets were stratified into training (80%) and test (20%). The model was tuned by further dividing the training set into training (80%) and validation (20%). The resulting trained model was evaluated on the test set. Robustness is evaluated by testing on an independent study cohort. **c)** Distribution of survival times in the study cohort. **d)** Top-10 features from Model-C as computed by SHAP. **e)** Annotation-free radiomics feature extraction pipeline. Dashed boxes indicate currently implemented methods in the pipeline where other options exist (e.g., sampling from pre-specified sites from an anatomical atlas as shown). ALP, alkaline phosphatase; BLTUMSZ, baseline tumor size; CL, chlorine; GGT, gamma glutamyl transferase; HCT, hematocrit; LDH, lactate dehydrogenase; MG, magnesium; NEUT, neutrophils; PD-L1, programmed death ligand 1; SHAP, SHapley Additive exPlanations; TSH, thyroid stimulating hormone.

Table 2 shows the stratification of patients in the different cohorts. The patient demographics along with the baseline tumor size are included in Table 2 for each cohort. With a median survival of 332 days (see Figure 1c) in the primary cohort, 1-year survival was selected as the outcome for modeling.

### 2.2 Feature Extraction

Pretreatment clinical-radiological information collected include patient demographics, routine blood work, and high-resolution CT imaging. The process of extracting features from both these modalities has been described below.

#### 2.2.1 Clinical Features

The clinical features available included demographic and blood work values. Categorical demographic information such as gender, smoking history, Eastern Cooperative Oncology Group (ECOG) performance status and so on were encoded into binary variables. Other numerical features collected were body mass index (BMI), blood work values such count of neutrophils, count of lymphocytes, amount of albumin, gamma-glutamyl transferase (GGT), alkaline phosphatase (ALP), and other available baseline models. Features that were highly correlated with each other were dropped from the analysis. The complete list of included clinical parameters and their units can be found in supplementary material (see Supplementary Table 3). Figure 1d shows the top features selected in the clinical model (Model-C).

#### 2.2.2 Whole-Lung Radiomic Features

CT images of the lung region were captured for each patient at baseline using whole-body, thoracic, or chest and abdominal CT imaging. Our in-house annotation tool was used to delineate the lung extents along the coronal and axial views in the abdominal and whole-body CT images, and the images were cropped to include only this lung region. Since the images were captured from multiple sites using several different scanners, intensity harmonization was performed by clipping CT gray values in the range of [-120–300] Hounsfield Units to focus on lung tissues (13). This was then followed by sampling patches from the image. Different sampling strategies can be employed such as anatomically informed sampling (based on lobes, nodes, etc.), uniform sampling, etc. We uniformly sampled the entire lung region and extracted n=80 3D patches of size 5cm^3^, as illustrated in Figure 1*e*. Finally, radiomic features such as statistical radiomic features and texture features including gray-level cooccurrence matrix (GLCM), gray-level run length matrix (GLRLM), gray-level difference matrix (GLDM), gray-level size zone (GLSZM), and neighborhood gray tone difference matrix (NGTDM) were extracted using PyRadiomics (14) with the 3D patches serving as the region of interest. The above proposed pipeline is novel and is shown in Figure 1*e*.

### 2.3 Machine Learning Modeling

The proposed whole-lung radiomics approach yielded radiomic features from a fixed number of patches in a given CT image. These features were then aggregated using simple summary statistics across the patches, namely mean, median, minimum, maximum, sum, mean absolute deviation, skew, and kurtosis, to provide a holistic view of the lung as well as to reduce the dimensionality problem that would otherwise arise. Feature selection was performed on the aggregated features on the training set using correlation and mutual information approaches with bootstrapping to further reduce the feature dimension. Specifically, from each pair of features with absolute Spearman’s correlation >0.9 a single feature is retained. Features with the highest mutual information with the target (OS) were selected. The process was repeated five times on separate bootstraps of the training set and only features selected at least twice made the final feature set. This was performed independently for the clinical and radiomic features. The selected features were then used as input to train an XGBoost (15) model. The parameters of the model were tuned using the training data and the threshold for prediction was determined using the receiver operating characteristics curve on the validation data. The final model was then generated by finetuning on the validation data^3^. This final model was used to evaluate the performance on the test set as well as the generalizability to an independent study cohort. SHapley Additive exPlanations (SHAP) (16) were used to identify important features contributing to the model performance as shown in Figure 1d. The overall analysis pipeline is shown in Supplementary Figure 4. Python (17) programming language was used for the feature extraction and selection pipelines. PyRadiomics (14) was used for feature extraction. The modeling and evaluation were implemented in R programming language (18). The final model parameters are shown in Supplementary Table 4. Cox Proportional Hazards (19) (Cox PH) was used to compute the hazard ratios in the evaluation and the KM method was used for survival analysis.

## 3 Results

We evaluated individual models using clinical features (Model-C), radiomic features (Model-R), and the combined clinical-radiomic model (Model-C+R). The following sections describe the survival risk stratification experiments and robustness experiments.

### 3.1 Survival-based Risk Stratification

Following the approach detailed in the Machine Learning modeling, tuned XGBoost models were evaluated on the test set. First Model-C, trained using the selected clinical parameters was evaluated on the held-out test set and an accuracy of 63% was achieved. Survival curves were generated using KM analysis and the patient stratification was quantified via the *p*-value. The clinical model achieved a significant *p*-value (0.00011) suggesting that it is capable of stratifying high- and low-risk patients on IO.

Next, the radiomics features only model (Model-R) was evaluated on the held-out test set and an accuracy of 54% was achieved which was much lower than that of the clinical model. The KM analysis similarly shows a non-significant separation with a *p*-value of 0.091. However, the separation in median survival times between low-risk and high-risk patients is 243 days, similar to that found in the PD-L1 status KM curves as shown in Figure 2b.

**Figure 2.**
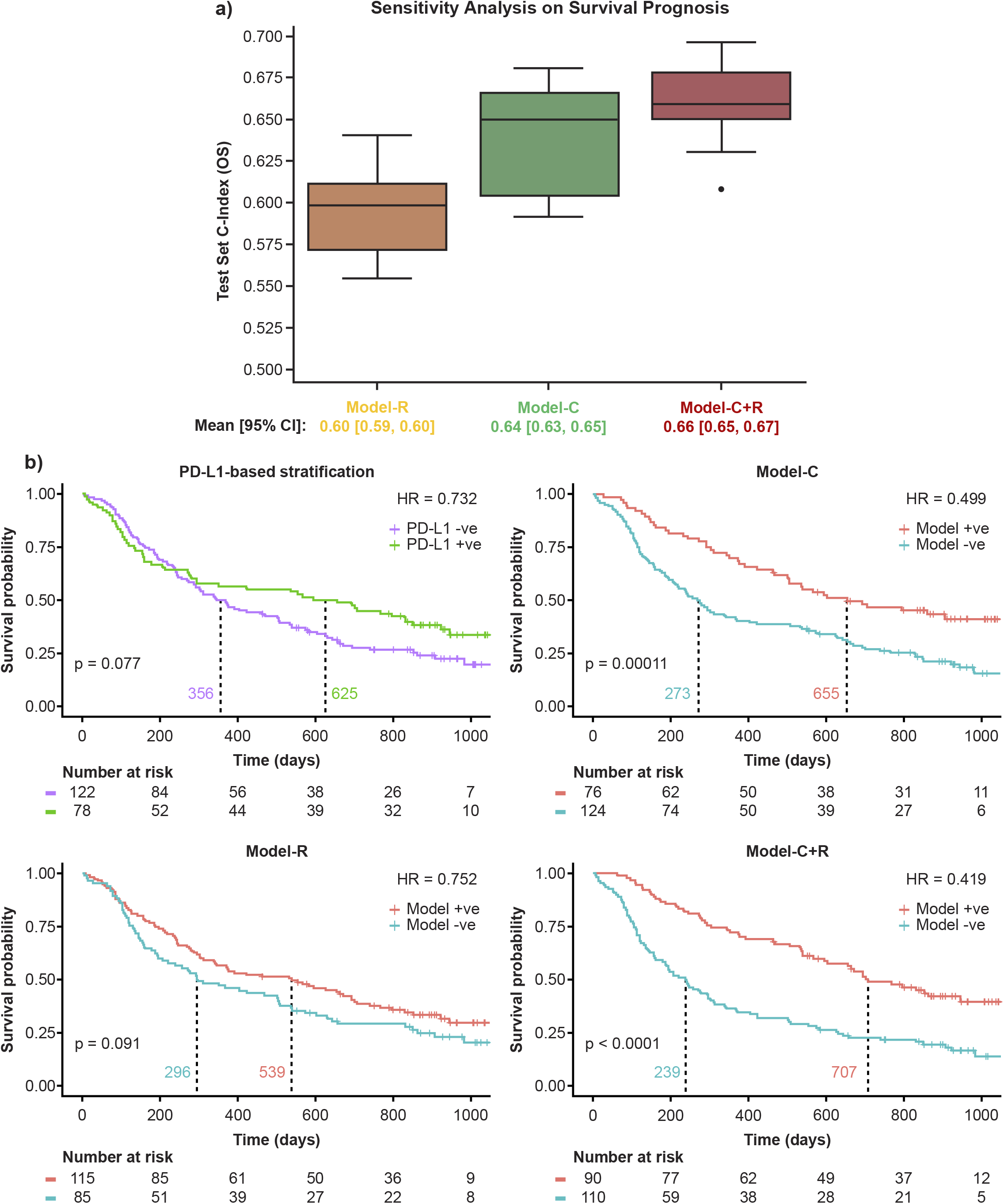
Model performance on the primary study cohort. Model+ve indicates model prediction of OS ≥ and Model-ve indicates model prediction of OS <365. **a)** Sensitivity analysis using 10 random train/test splits and **b)** Kaplan-Meier curves of the model performance on the held-out test set. +ve, positive; -ve, negative; HR, hazard ratio; PD-L1, programmed death ligand 1.

We then evaluated the potential complementary nature of the clinical and radiomic features using Model-C+R. The combined model yielded an accuracy of 68% on the held-out test set, a 5% gain over Model-C. The SHAP feature importance plot for the model was generated and the importance and contribution of radiomic features alongside clinical ones was noted in Supplementary Figure 6 (top panel). The KM analysis yielded a *p*-value <0.0001 suggesting a significant separation of high- and low-risk patients. Furthermore, separation in median survival times was largest using Model-C+R, at 486 days.

The large separation in median survival times suggests the benefit of combining radiomic and clinical models. The KM curves for the models are shown in Figure 2b.

### 3.2 Model Robustness

#### 3.2.1 Sensitivity to Data Splits

Machine learning algorithms are often affected by potential unseen biases in the selection of the training/test splits. To test the sensitivity of the selected features to the selection of training/test splits, 10 random training/test splits were tested on the OS prediction task. The results, shown in Figure 2a, validate the stability of the selected features to training/test splits with consistent concordance indices and a tight 95% confidence interval across all models.

#### 3.2.2 Generalization to Independent Study Cohort

The independent study cohort was used to validate the models trained on the training set. The survival distribution of the independent study cohort, shown in Figure 3a, had a median survival of 282 days, 35 days lower than that of the training set (primary study cohort). Previously selected features were extracted from the independent study cohort and tested using the three models: Model-C, Model-R, and Model C+R trained on the primary study cohort training data. Similar to the performance observed on the test set (primary study cohort), Model-C outperformed Model-R, and Model-C+R outperformed both the clinical and radiomic models, further confirming the complementary nature of the clinical and radiomic features. The model achieved an accuracy of ∼72% and stratified the patients into high- and low-risk groups with a difference in median survival of 423 days as shown in Figure 3b. The comparison of the performance of the different models across both test cohort and independent study cohorts in terms of sensitivity, specificity, precision and accuracy are reported in Supplementary Table 6. SHAP feature importance plot of Model-C+R on the independent study cohort revealed similar top features with the same radiomic features contributing to the prediction (Supplementary Figure 6, bottom panel). Further details on the SHAP analyses as well as the building of more traditional Cox models using the top predictors of overall survival are included in the Supplementary Material – SHAP Analysis.

**Figure 3.**
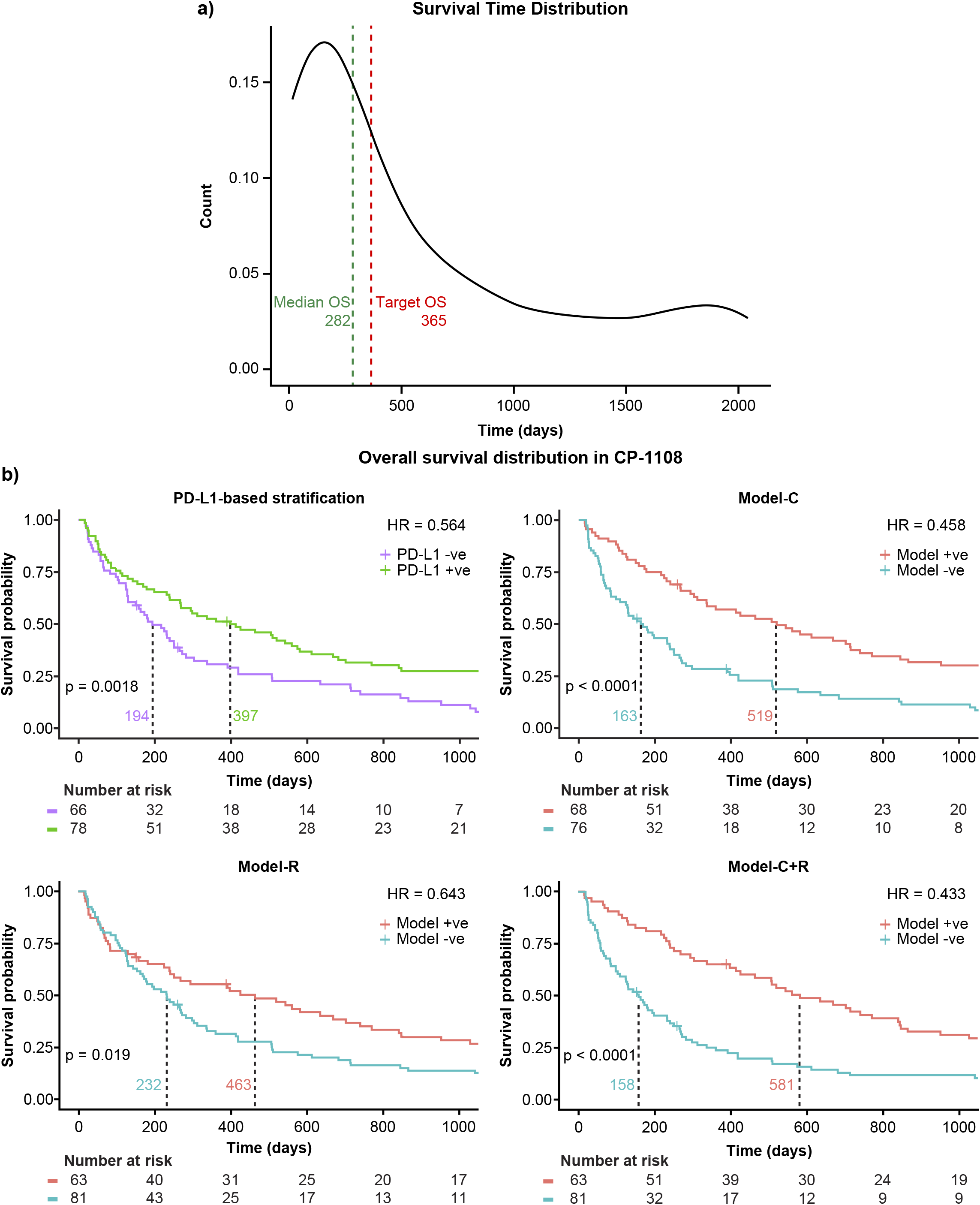
Generalization to an independent study cohort. Model+ve indicates model prediction of OS≥365 and Model-ve indicates model prediction of OS <365. **a)** Survival distribution in the independent study cohort and **b)** Kaplan-Meier curves of the model performance on the independent study cohort. +ve, positive; -ve, negative; HR, hazard ratio; PD-L1, programmed death ligand 1.

## 4 Discussion

The emergence of PD-L1 inhibitors as first line treatment for non-resectable advanced NSCLC^2^ has provided improved quality of life and an increased survival time. In this paper we have presented a novel approach to the prediction of patients’ survival that integrates clinical and radiomic features and requires no previous tumor annotation. Our approach is competitive in terms of concordance index (when predicting overall survival) and accuracy (when predicting risk of less than median survival). Finally, our approach compares very favorably with predictions based on PD-L1 status and has been validated against a multi-trial test set as well as an independent additional trial.

The potential for such an approach as a diagnostic tool is explored by including the standard of care (SoC) arms from the clinical trials of MYSTIC and NEPTUNE^4^. The KM curves from the model predictions are illustrated in Figure 4. Model-C+R offers the potential to select patients who would benefit from IO therapy. As shown in Figure 4a, Model-C+R identifies patients who would not benefit from IO and have similar survival curves to the SoC arms. It also identifies patients who would benefit from IO therapy and when combined with PD-L1 expression helps identify patients with the highest odds of survival (Figure 4b). The Model-C+R identified patients that also satisfy a higher (50%) cutoff for PD-L1 show the best improvement in median OS as shown in Figure 4d. Thus, the proposed non-invasive approach opens new avenues for patient selection that are complementary to PD-L1 in the mNSCLC setting.

**Figure 4.**
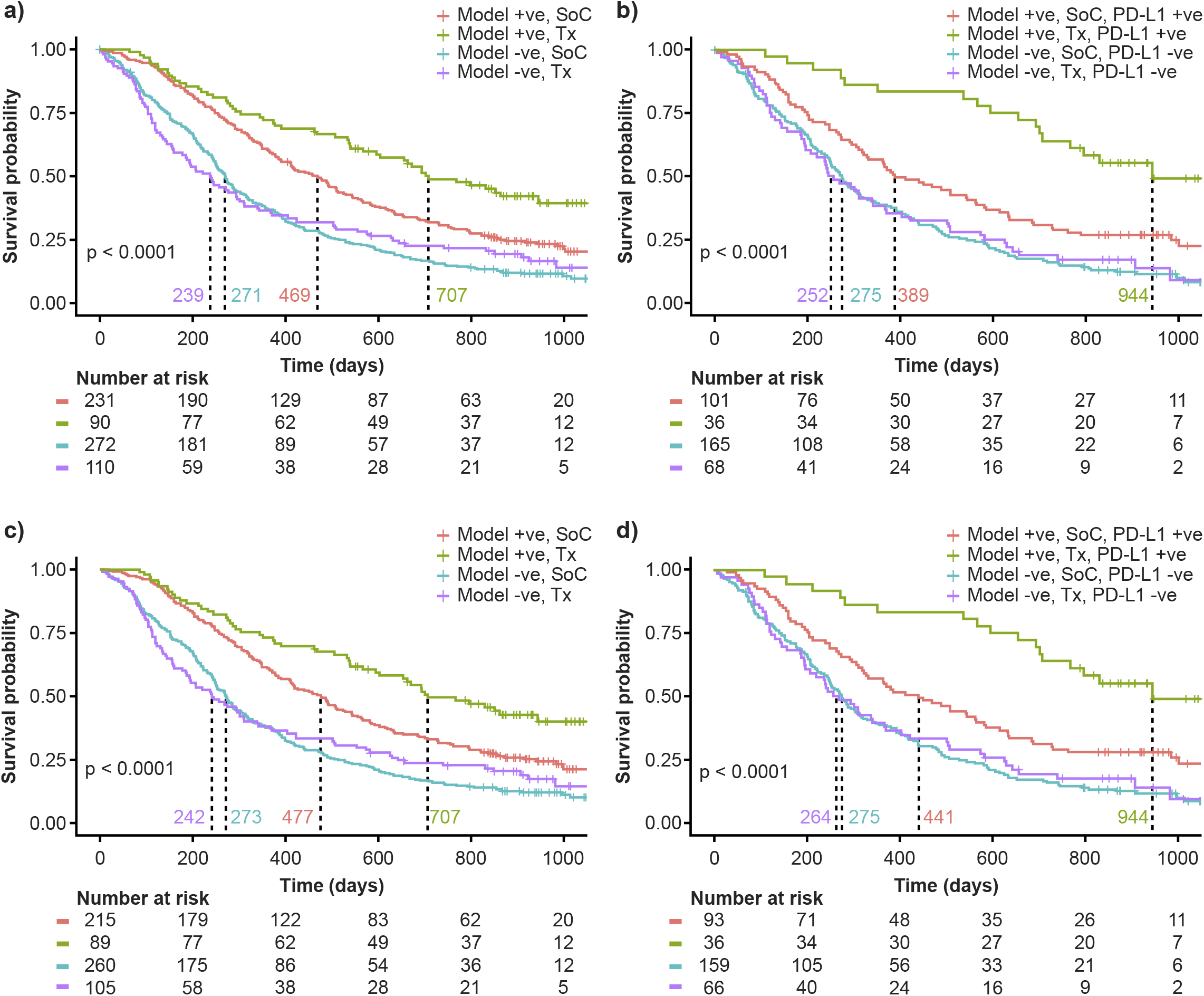
Evaluating the potential of Model-C+R as a diagnostic tool for survival-based patient risk stratification. Model+ve indicates model prediction of OS≥365 and Model-ve indicates model prediction of OS <365. **a)** Comparing model risk stratification on Tx arm against SoC. **b)** Survival risk stratification on PD-L1 status and model prediction intersection in the test set (primary study cohort) and SoC. Patients identified as low-risk on Tx have median survival 944 days, 692 days more than high-risk patients on Tx. **c)** Comparing model risk stratification on Tx arm against SoC for sub-cohort with PD-L1 expression 50% cutoff available. **d)** Survival risk stratification on PD-L1 status at 50% cutoff and model prediction intersection in the test set (primary study cohort) and SoC. Patients identified as low-risk on Tx have median survival of 944 days. +ve, positive; -ve, negative; PD-L1, programmed death ligand 1; SoC, standard of care; Tx, treatment.

### 4.1 Challenges and Limitations

In order to build clinician confidence around using such diagnostic tools, it is important to build an intuitive understanding of the interpretability of these features. The radiomics features identified by our models to be most informative are shown in the Supplementary Material – SHAP Analysis. Our models suggest that low texture heterogeneity corresponds to better OS. This is similar to results reported in literature where texture features representing tumor heterogeneity correlated with OS and PFS (20). However, with the exploration of the whole lung, these features need to be further investigated in terms of interpretability and consistency with the biology of cancer.

When the clinical trials were originally designed, the optimal cutoff for PD-L1 expression in first line mNSCLC was not well known; consequently, while valid based on data at the time, we have only 25% cutoff available consistently for PD-L1 in the studies explored herein. It is now more broadly known that 50% cutoff is more clinically relevant for IO monotherapy. The performance of Model-C+R on the patient subset with 50% PD-L1 cutoff is shown in Supplementary Figure 5 and further analysis with additional datasets will be performed to establish clinical relevance and clinician confidence.

The ML models presented in this work were developed using only data from IO treatment arms. This makes it challenging to tease out the predictive/prognostic nature of the features used by the model. We plan to address this limitation as a key next step in our future research.

Though we have shown generalizability of our model to a completely independent study cohort, prospective validation of such models will be the key towards the development of diagnostic tools that can be used in the clinic.

### 4.2 Future Work

An important next step is to identify the predictive and prognostic nature of the model features to assist in development of further diagnostic tools. Additionally, we also plan to explore the possibility of using imaging data from follow-ups (*delta*-radiomics) to further improve our models. Results in these directions will be reported elsewhere.

## 5 Conflict of Interest

Kedar Patwardhan, Harish RaviPrakash, Domingo Salazar and Paul Metcalfe are employees of AstraZeneca. Ignacio Gonzalez-Garcia, Joachim Reischl, Nikos Nikolaou were employees of AstraZeneca at the time of contribution to the manuscript.

## 6 Author Contributions

- **KP** – Conceptualization, Methodology, Software, Validation, Formal Analysis, Investigation, Data Curation, Writing – Original Draft, Writing – Review & Editing, Supervision, Project Administration
- **HR** – Methodology, Software, Validation, Formal Analysis, Investigation, Data Curation, Writing – Original Draft, Writing – Review & Editing, Visualization
- **NN** – Methodology, Software, Validation, Formal Analysis, Investigation, Writing – Original Draft, Writing – Review & Editing, Visualization
- **IG** – Methodology, Investigation, Data Curation, Writing – Original Draft, Writing – Review & Editing
- **DS** – Methodology, Software, Validation, Formal Analysis, Investigation, Writing – Original Draft, Writing – Review & Editing
- **PM** – Resources, Writing – Review & Editing
- **JR** - Conceptualization, Resources, Writing - Review & Editing, Supervision, Funding acquisition

## 7 Funding

The work done was supported by AstraZeneca.

## 8 Acknowledgments

The authors would like to thank Paul Agapow and Evan Wu for useful inputs, technical discussions and annotations.

## Footnotes

1 For additional details regarding the selection of these specific references, please refer to the exact search link^4^ and Supplementary Material - Literature Review.

2 The data for 25% cutoff is consistently available across all the studies used for our analysis. KM curves for different sub-cohorts where we may have 50% cutoff information available are provided in Supplementary Figure 3.

3 Further details on the training of the XGBoost model can be found in Supplementary Material – *Analysis Pipeline*.

4 Patient Demographics from the SoC arms can be found in Supplementary Material – Data Preparation.

## Notes

### Competing Interest Statement

All author contributions were made when the authors were employed by AstraZeneca. There are no other competing interests to be disclosed.

## References

1. Cancer.Net. Lung Cancer - Non-Small Cell: Statistics. 2022.

2. Paz-Ares L, Spira A, Raben D, Planchard D, Cho B, Özgüroğlu M, et al. Outcomes with durvalumab by tumour PD-L1 expression in unresectable, stage III non-small-cell lung cancer in the PACIFIC trial. 2020;31(6):798–806.

3. Rizvi NA, Cho BC, Reinmuth N, Lee KH, Luft A, Ahn M-J, et al. Durvalumab with or without tremelimumab vs standard chemotherapy in first-line treatment of metastatic non–small cell lung cancer: the MYSTIC phase 3 randomized clinical trial. 2020;6(5):661–74.

4. PubMed. https://pubmed.ncbi.nlm.nih.gov/?term=%22nsclc%22%20and%20%22immunotherapy%22%20and%20%22radiomics%22%20and%20%22CT%22%20and%20%22advanced%22%20not%20%22PET%22&filter=simsearch1.fha&sort=date.

5. ClinicalTrials.gov. A Global Study to Assess the Effects of MEDI4736 (Durvalumab), Given as Monotherapy or in Combination With Tremelimumab Determined by PD-L1 Expression Versus Standard of Care in Patients With Locally Advanced or Metastatic Non Small Cell Lung Cancer (ARCTIC).

6. Planchard D, Reinmuth N, Orlov S, Fischer J, Sugawara S, Mandziuk S, et al. ARCTIC: durvalumab with or without tremelimumab as third-line or later treatment of metastatic non-small-cell lung cancer. 2020;31(5):609–18.

7. ClinicalTrials.gov. Phase III Open Label First Line Therapy Study of MEDI 4736 (Durvalumab) With or Without Tremelimumab Versus SOC in Non Small-Cell Lung Cancer (NSCLC).

8. ClinicalTrials.gov. Study of Durvalumab With Tremelimumab Versus SoC as 1st Line Therapy in Metastatic Non Small-Cell Lung Cancer (NSCLC) (NEPTUNE).

9. de Castro Jr G, Rizvi NA, Schmid P, Syrigos K, Martin C, Yamamoto N, et al. NEPTUNE: phase 3 study of first-line Durvalumab plus tremelimumab in patients with metastatic NSCLC. 2023;18(1):106–19.

10. Kaplan EL, Meier PJJotAsa. Nonparametric estimation from incomplete observations. 1958;53(282):457–81.

11. Johnson ML, Cho BC, Luft A, Alatorre-Alexander J, Geater SL, Laktionov K, et al. Durvalumab with or without tremelimumab in combination with chemotherapy as first-line therapy for metastatic non–small-cell lung cancer: the phase III POSEIDON study. 2023;41(6):1213.

12. ClinicalTrials.gov. A Phase 1/2 Study to Evaluate MEDI4736.

13. Trebeschi S, Bodalal Z, Boellaard TN, Tareco Bucho TM, Drago SG, Kurilova I, et al. Prognostic value of deep learning-mediated treatment monitoring in lung cancer patients receiving immunotherapy. 2021:9.

14. Van Griethuysen JJ, Fedorov A, Parmar C, Hosny A, Aucoin N, Narayan V, et al. Computational radiomics system to decode the radiographic phenotype. 2017;77(21):e104–e7.

15. Chen T, He T, Benesty M, Khotilovich V, Tang Y, Cho H, et al. Xgboost: extreme gradient boosting. 2015;1(4):1–4.

16. Lundberg SM, Lee S-IJAinips. A unified approach to interpreting model predictions. 2017;30.

17. vanRossum GJDoCS. Python reference manual. 1995(R 9525).

18. Team RC. R: A language and environment for statistical computing. 2013.

19. Cox DR. Regression models and life tables. Journal of the Royal Statistical Society: Series 1 . L B. 1972;34(2):187–202.

20. Jazieh K, Khorrami M, Saad A, Gad M, Gupta A, Patil P, et al. Novel imaging biomarkers predict outcomes in stage III unresectable non-small cell lung cancer treated with chemoradiation and durvalumab. 2022;10(3).

21. Gong J, Bao X, Wang T, Liu J, Peng W, Shi J, et al. A short-term follow-up CT based radiomics approach to predict response to immunotherapy in advanced non-small-cell lung cancer. 2022;11(1):2028962.

22. He B-X, Zhong Y-F, Zhu Y-B, Deng J-J, Fang M-J, She Y-L, et al. Deep learning for predicting immunotherapeutic efficacy in advanced non-small cell lung cancer patients: a retrospective study combining progression-free survival risk and overall survival risk. 2022;11(4):670.

23. Liu C, Gong J, Yu H, Liu Q, Wang S, Wang JJFiO. A CT-based radiomics approach to predict nivolumab response in advanced non-small-cell lung cancer. 2021:15.

24. Trebeschi S, Drago S, Birkbak N, Kurilova I, Călin A, Pizzi AD, et al. Predicting response to cancer immunotherapy using noninvasive radiomic biomarkers. 2019;30(6):998–1004.

