## Supplementary Material for "Towards a Survival Risk Prediction Model for Metastatic NSCLC Patients on Durvalumab Using Whole-Lung CT Radiomics"

### 1 Literature Review

Supplementary Table 1. Overview of Prior Work Not Relevant to OS Prediction Using Baseline CT Images in Patients on IO, in a mNSCLC Setting.

| **Reference** | **Total no. of Patients** | **Test Cohort Size** | **AUC in Test Cohort** | **Target** |
| --- | --- | --- | --- | --- |
| Shen et al (1) | 42 | N/A | 0.80 (0.67–0.930 | PFS |
| Wen et al (2) | 120 | 30 | 0.79 (0.71–0.87) | PD-L1 |
| Yang et al (3) | 200 | 40 | 0.80 (0.74–0.86) | OR |
| Sun et al (4) | 390 | 130 | 0.848 (0.48–0.79) | PD-L1 |
| Vaidya et al (5) | 109 | 79 | 0.76 (N/A) | Hyperprogressors |
| Liu et al (6) | 161 | 49 | 0.71 (0.65–0.78) | OR |

Values not reported are indicated with “N/A” (i.e. not applicable).

IO, immunotherapy; mNSCLC, metastatic non-small cell lung cancer; OR, overall response rate; OS, overall survival; PD-L1, programmed death ligand 1; PFS, progression-free survival

Yang et al (7) combined radiomics and independent risk factors to predict progression-free survival (PFS) (not overall survival [OS], which is the focus of our work) using multivariate Cox Regression. Sun et al (4) worked on programmed death ligand 1 (PD-L1) expression prediction using radiomics and clinical features and was thus not included in the comparison. Similarly, Wen et al (2) used radiomics for tumor burden and PD-L1 expression prediction. Shen et al1 worked on the problem of response prediction on a small dataset of 42 patients (article in Chinese) with accuracy of 80.4%. Yang et al (3) used deep learning with 5-fold cross-validation on a 200 patient dataset to predict clinical outcomes. However, no accuracy or C-index was reported (only area under the curve [AUC]). Vaidya et al (5) predicted hyperprogressors using radiomics on a dataset of 109 patients (79 for test). The model reported an accuracy of 83% (F1 of 0.58, imbalanced dataset). Liu et al (6) used delta radiomics for response prediction in a dataset of 161 patients and achieved a C-index of 0.81 (0.68–0.95) on the test set (N_test_ = 49). The radiomics model on baseline data performed poorly with an AUC of 0.61.

### 2 Data Preparation

The data from the treatment arms of the three clinical trials of MYSTIC, ARCTIC, and NEPTUNE are combined and Kaplan-Meier (KM) survival analysis was performed to observe the difference in survival with respect to clinical trials, type of treatment (mono vs combination) and PD-L1 expression. Supplementary Figure 2a shows the survival analysis based on the different clinical trials. The median survival in the three trials differs by a difference of at most 75 days (*p* > 0.05).

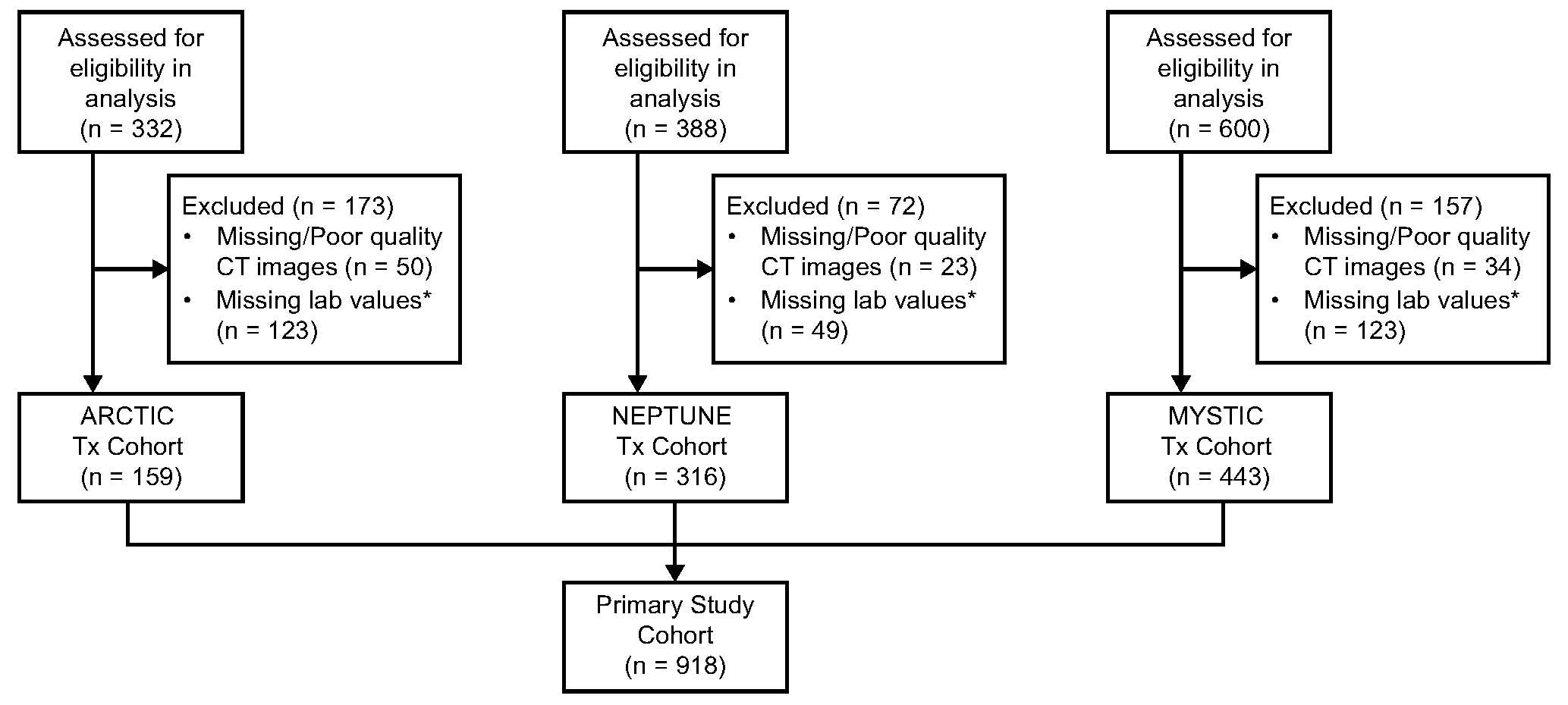

Supplementary Figure 1. Data selection for analysis in the treatment (Tx) arms of the clinical studies. *Lab values refers to the clinical parameters listed in Supplementary Table 3.

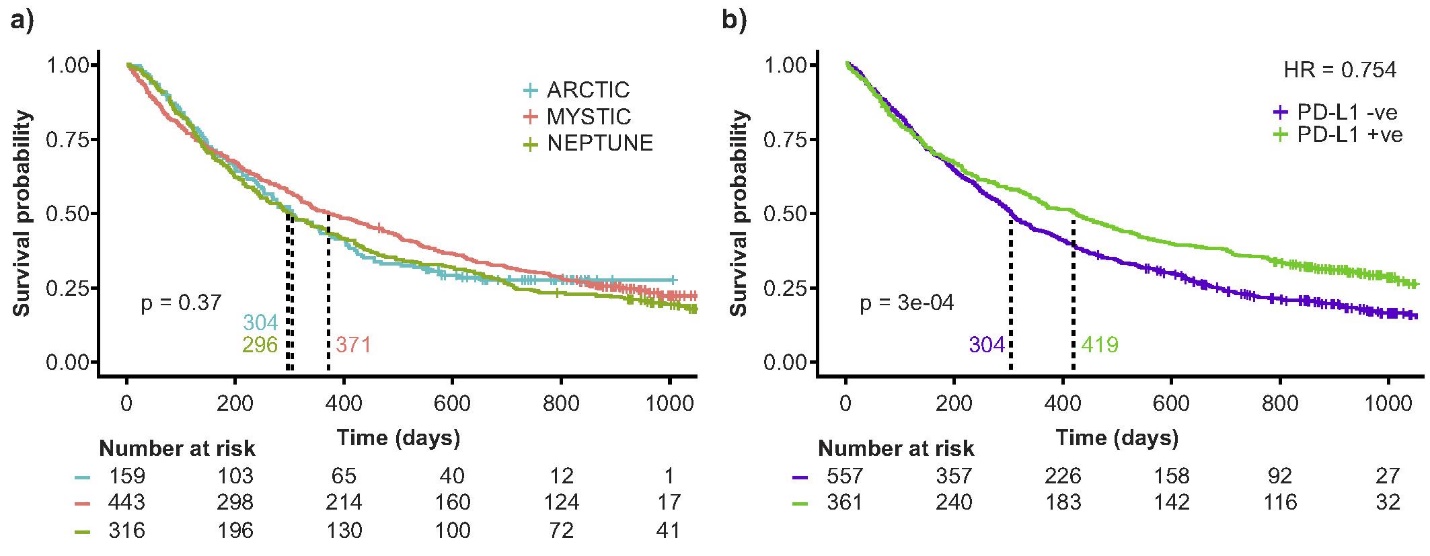

Supplementary Figure 2. KM curves of survival times based on PD-L1 stratification in the primary study cohort. a) KM curves based on different clinical studies and b) KM curves for PD-L1 expression based stratification. +ve, positive; -ve, negative; HR, hazard ratio; KM, Kaplan-Meier; PD-L1, programmed death ligand 1.

With durvalumab being a targeted checkpoint inhibitor, the survival distributions of patients with respect to PD-L1 cut-off was evaluated. The survival was observed to be significantly different with a *p*-value < 0.05 as shown in Supplementary Figure 2b.

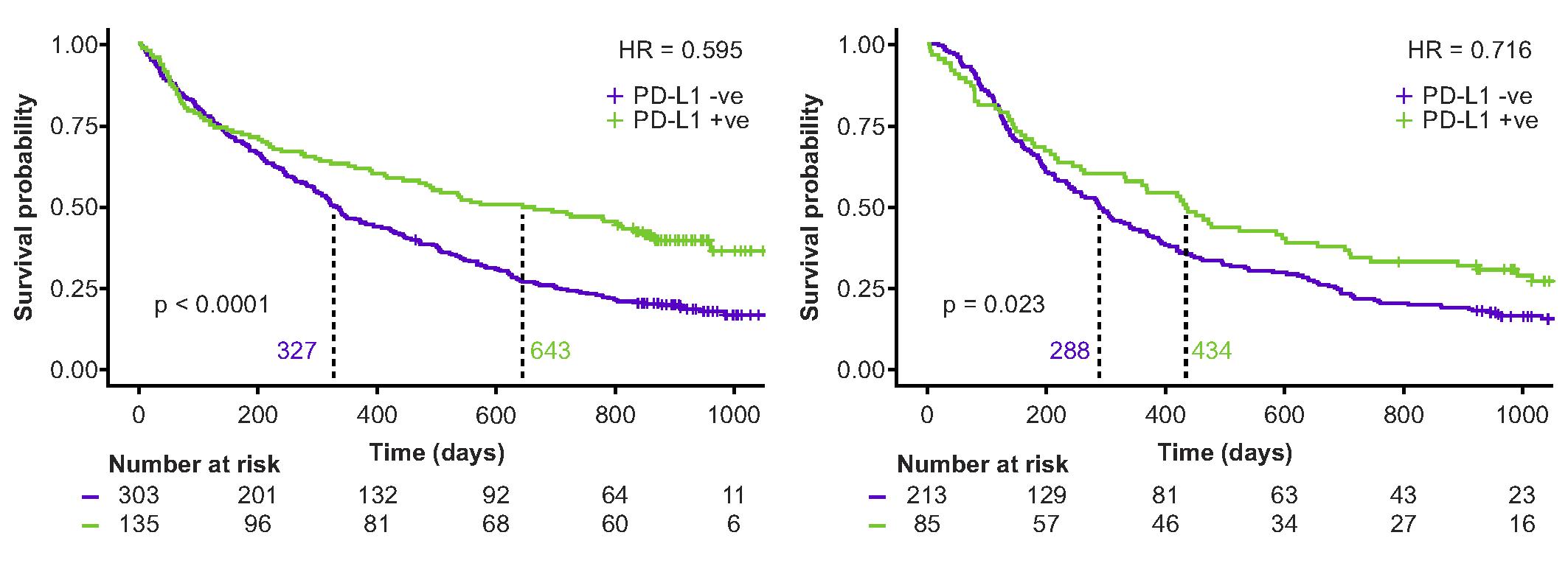

Supplementary Figure 3. PD-L1 distribution at 50% cutoff in primary sub-cohorts of MYSTIC (left) and NEPTUNE (right). +ve, positive; -ve, negative; HR, hazard ratio; PD-L1, programmed death ligand 1.

#### 2.1 Standard of Care Data

Standard of care (SoC) data from the three clinical trials were retrospectively assessed for eligibility to be included in the analysis. At the time of analysis, due to lack of clarity on the SoC data in the arctic clinical trial, this data was excluded. Thus, data from the MYSTIC and NEPTUNE trials were used and exclusion criteria of unevaluable OS, poor quality/missing radiological data at baseline, and missing clinical parameters was used and the final SoC cohort comprised of 503 patients. The patient demographics were compared to that of the test set and found to be predominantly similar as shown in Supplementary Table 2.

#### 2.2 Clinical Features

Routine blood work variables were collected at baseline such as lymphocyte count, neutrophil count, amount of gamma-glutamyl transferase enzyme in the blood, and many more. Variables with over 10% missingness in the data were excluded prior to evaluating the eligibility of patients for the analysis. Correlation of variables in the patients with evaluable OS was performed and highly correlated variables were excluded. A complete list of the included clinical parameters can be found in Supplementary Table 3. Patients with missing data in the identified parameters were also excluded from the analysis.

Supplementary Table 2. Patient Demographics in SoC Arms of MYSTIC and NEPTUNE Studies as Compared to the Test Set in the Primary Study Cohort.

| **Patient Cohorts** | | | | | | |
| --- | --- | --- | --- | --- | --- | --- |
| **Study Name** | **SoC** | | | **Test Set** | | |
|  | **PD-L1 Status** | | | **PD-L1 Status** | | |
|  | **Neg.** | **Pos.** | | **Neg.** | | **Pos.** |
| ARCTIC | - | - | | 29 | | 7 |
| MYSTIC | 128 | 79 | | 51 | | 43 |
| NEPTUNE | 167 | 129 | | 42 | | 28 |
| All data by PD-L1 status | 295 | 208 | | 122 | | 78 |
| All data combined | 503 | | | 200 | | |
| **Patient Demographics and Imaging Characteristics** | | | | | | |
| **Covariate** | | | **SoC** | | **Test Set** | |
| Sex = Female | | | 26% | | 27% | |
| Age (years): Mean (± SD) | | | 63.19 (± 8.97) | | 62.9 (± 9.99) | |
| Overall survival: Median (IQR) | | | 345 (190–668) | | 385 (149–836) | |
| Baseline tumor size: Median (IQR) | | | 77 (49–121) | | 83 (50–121) | |
| In-plane resolution (mm): Mean (± SD) | | | 0.78 (± 0.10) | | 0.78 (± 0.10) | |
| Slice thickness (mm): Mean (± SD) | | | 4.3 (± 1.05) | | 4.4 (± 1.24) | |
| CT scanner manufacturers (Top – 4) | | | SIEMENS (34%) GE (31%) TOSHIBA (16%) PHILIPS (17%) | | GE (33%) SIEMENS (31%)  PHILIPS (20%) TOSHIBA (15%) | |

IQR, interquartile range; Neg., negative; PD-L1, programmed death ligand 1; Pos., positive; SD, standard deviation

Supplementary Table 3. Table of Clinical Parameters Extracted From Blood Tests and Demographic Information Used in the Development of Model-C. Units of Each Parameter is Indicated Below the Respective Parameters.

| **Clinical Parameters** | | | | |
| --- | --- | --- | --- | --- |
| Chloride (mmol/L) | Hematocrit | Calcium (mmol/L) | Baseline BMI (kg/m^2^) | Combined amount of both albumin and globulin |
| Albumin (g/L) | Baseline Tumor Size (mm) | Alkaline Phosphatase (U/L) | Glucose (mmol/L) | Magnesium (mmol/L) |
| Monocytes (10^9^/L) | Neutrophils (10^9^/L) | Platelets (10^9^/L) | Potassium (mmol/L) | Treatment Combination |
| Lactate Dehydrogenase (U/L) | Gamma-Glutamyl Transferase (U/L) | Eosinophil (10^9^/L) | Basophils (10^9^/L) | Sex |
| Thyroid Stimulating Hormone | Lymphocytes (10^9^/L) | Aspartate amino Transferase (U/L) | Smoking History | Baseline WHO Performance Status |

BMI, body mass index; WHO, World Health Organization.

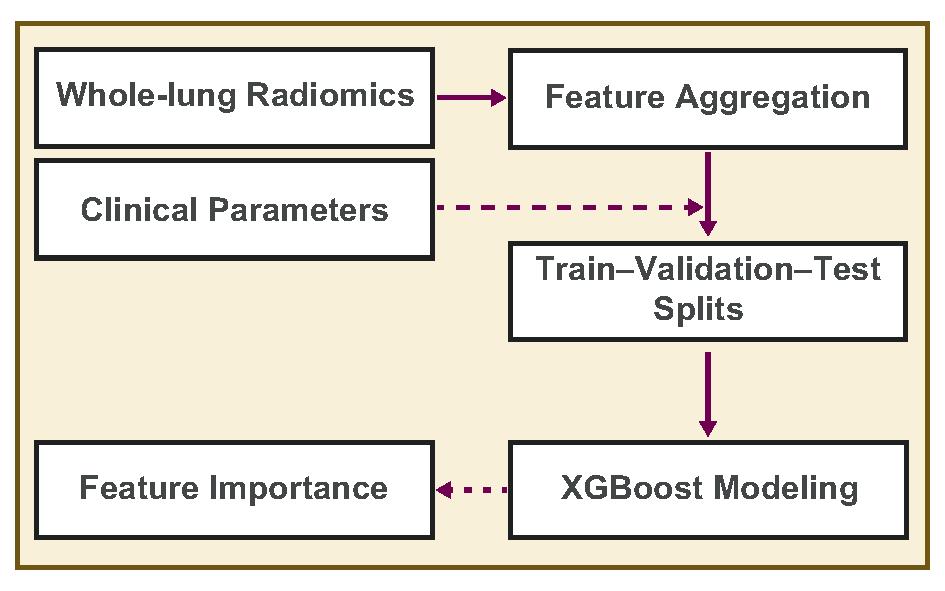

Supplementary Figure 4. Analysis pipeline starting with the feature extraction step, followed by feature aggregation (across patches), addition of clinical parameters, splitting of data for modeling, training of XGBoost model and ending with the feature importance.

#### 2.3 Analysis Pipeline

The overall analysis pipeline is shown in Supplementary Figure 4. The whole-lung radiomic features extracted from each patch were aggregated using summary statistics such as mean, median, minimum, maximum, skew, kurtosis, mean absolute deviation, and sum. These eight statistics were computed for each radiomic feature for each patient. Depending on the modeling approach, clinical parameters were appended to the radiomic feature set. The data was then split into predefined train-validation-test cohorts which were used to train XGBoost models. Feature importance was evaluated using Shapley values to interpret the contribution of the different modalities and features.

The steps involved in the survival-based risk stratification and generalization experiments are described in detail below.

##### 2.3.1 Survival-Based Risk Stratification Modeling Step

XGBoost Tree model was tuned using the *mlr* package in R. 5-fold stratified cross-validation was used to tune the model parameters of depth (max_depth), child weight (min_child_weight), subsampling ratio of training instances (subsample), and subsample ratio of columns in constructing trees (colsample_bytree) with binary logistic objective function and learning rate of 0.001. The tuned parameters are listed in Supplementary Table 4.

Supplementary Table 4. Final Model Parameters for Model-C, Model-R, and Model-C+R After Tuning.

| Parameters | Model-C | Model-R | Model-C+R |
| --- | --- | --- | --- |
| Max_depth | 6 | 7 | 4 |
| Min_child_weight | 2.01866 | 9.917206 | 2.643712 |
| Subsample | 0.6441625 | 0.8392963 | 0.8810002 |
| Colsample_bytree | 0.9481753 | 0.5950314 | 0.6997333 |

Following this tuning step, the model was trained with the training set for up to 5000 iterations and stopped when validation set loss stopped decreasing for 100 iterations. Receiver operating characteristic (ROC) curve for prediction on validation set was generated for the above trained model. A threshold for prediction was identified as the value at which the product of sensitivity and specificity was highest (so as to not bias the model towards either metric). The model was then finetuned with the validation data to generate the final model. This final model and the previously identified threshold were used to predict survival risk on the unseen test set.

##### 2.3.2 Sensitivity to Data Splits Modeling Step

80% of the training data (selected uniformly at random) were used for training five different survival models for predicting OS. The model list includes two linear: Cox PH with Ridge (L2-norm) regularization and Cox PH with Elastic Net (L2-norm and L1-norm) regularization as well as three non-linear ones: Survival Forest, Gradient Boosted Cox PH, Gradient Boosted Component-wise Least Squares. The remaining 20% of the training data was used to finetune the model hyperparameters. Once the hyperparameters were optimized, their values were fixed and the five models were retrained on the full training set. A heterogeneous ensemble of the five models was used for the final predictions on the test set. The OS ranking prediction on each patient was given by a linear combination of the individual predictions of the five models (each normalized within the range [0–1] and the weight of the i^th^ model being equal to w_i_ = C-index_i_ – 0.5 (its ‘edge’ over random survival prediction) as measured on the training set. Future improvements could include the weighs w_i_ being based on a separate validation set rather than the full training set. For evaluation purposes we report the ensemble’s C-index measured on the test set.

Supplementary Table 5. List of Metrics Used in the Evaluation of the Models.

| **Metric** | **Definition** |
| --- | --- |
| Accuracy | Ratio of correctly identified high-risk and low-risk patients to the complete pool of patients |
| Sensitivity | Ratio of correctly identified high-risk patients to all who are high-risk patients in reality |
| Specificity | Ratio of correctly identified low-risk patients to all who are low-risk patients in reality |
| Precision | Ratio of correctly identified high-risk patients to all identified high-risk patients |
| Area Under the Receiver Operating Characteristics Curve (AUC) | Defines the proportion of high-risk patients ranked before a uniformly drawn random low-risk patient |
| Concordance Index | Defines the fraction of pairs in the data where the observation with higher survival time has higher probability of survival predicted by the model |

AUC, area under the curve.

##### 2.3.3 Sensitivity Analysis Step

For the sensitivity analysis, the entire process of splitting the data at random into a training and test set, performing feature selection and training, and OS evaluating models (as described above) was repeated 10 times to investigate the performance of training models under each modality (clinical, radiomics, clinical + radiomics) on different train/test splits. We reported the average C-index measured on the test set and its 95% confidence interval, as well as its median value and interquartile range (IQR).

### 3 Results

#### 3.1 Metrics Used For Evaluation

Metrics used for evaluation of the risk stratification and sensitivity analysis are discussed in Supplementary Table 5. Sensitivity and specificity are metrics that may be used in a clinical setting by physicians to compare different patient stratification models.

Supplementary Table 6. Model Performance on Test cohort and Independent study cohort.

| Model | Sensitivity | Specificity | Precision | Accuracy |
| --- | --- | --- | --- | --- |
| **Test Cohort** | | | | |
| Model-C | 0. 505 | **0.758** | 0.697 | 0.631 |
| Model-R | **0.619** | 0.474 | 0.565 | 0.546 |
| Model-C+R | **0.619** | 0.737 | **0.722** | **0.678** |
| **Independent Study Cohort** | | | | |
| Model-C | 0.644 | 0.647 | 0.559 | 0.646 |
| Model-R | 0.576 | 0.659 | 0.54 | 0.618 |
| Model-C+R | **0.695** | **0.741** | **0.651** | **0.718** |

#### 3.2 Model Performance

The performance of Model-C, Model-R, and Model-C+R on the test set and independent study cohort are tabulated in Supplementary Table 6. Model-C+R shows superior performance to both Model-C and Model-R in both cohorts. Independently Model-R and Model-C have complementary performance with respect to sensitivity and specificity, but Model C+R has the most balanced sensitivity and specificity across both test cohorts.

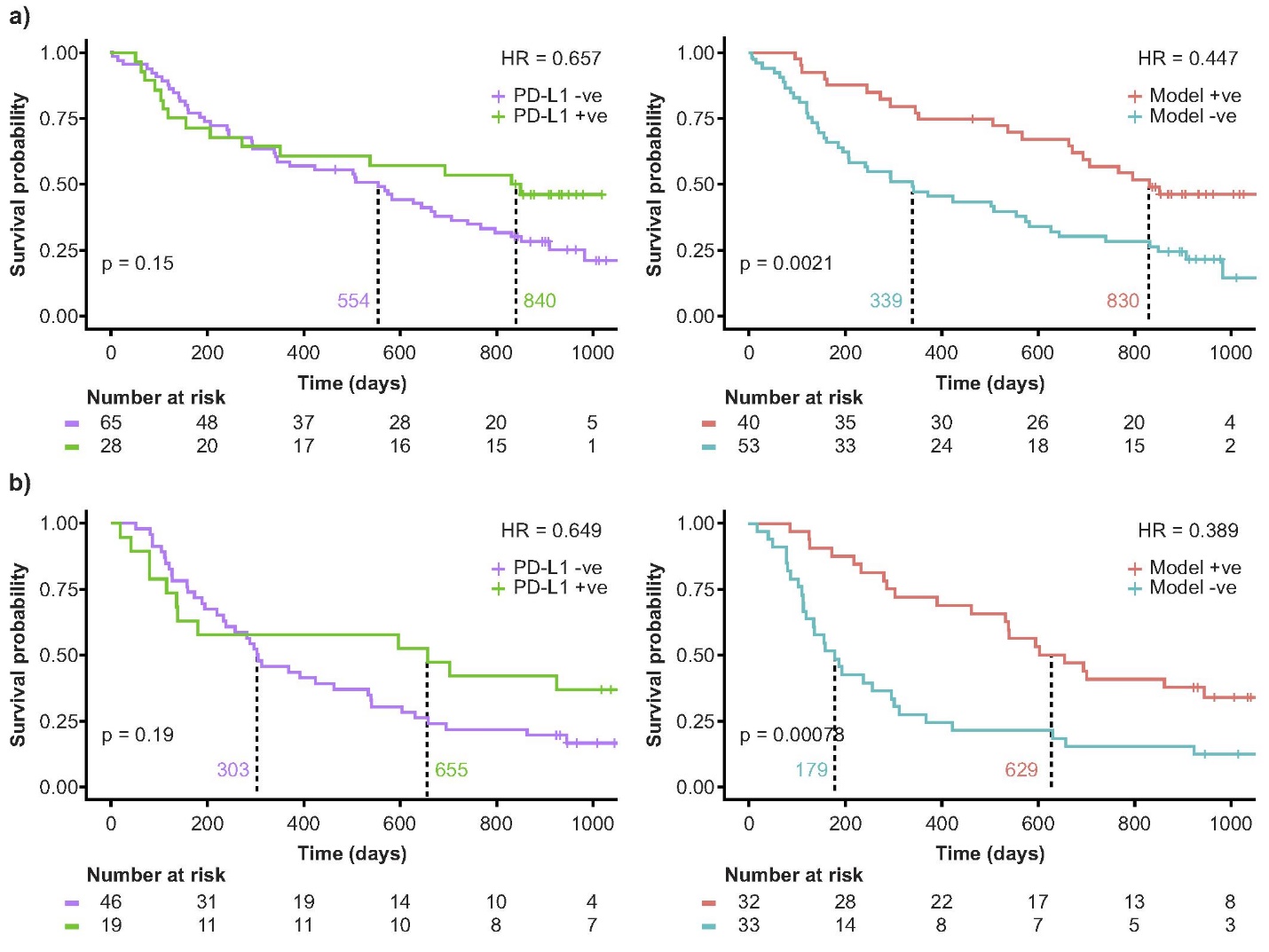

Supplementary Figure 5. Comparison of Kaplan-Meier curves of survival times of PD-L1 50% cutoff and Model-C+R (in the test cohort only). a) Comparison in MYSTIC study and b) comparison in NEPTUNE study. +ve, positive; -ve, negative; HR, hazard ratio; PD-L1, programmed death ligand 1.

#### 3.3 Model Performance vs PD-L1 at 50% Cutoff

With 50% tumor cells positive for PD-L1 expression as the FDA approved cutoff in other immunotherapy drugs, we compare the performance of Model-C+R against PD-L1 50% cutoffs wherever available. Particularly, this information was available for MYSTIC and NEPTUNE studies. The model performance was compared to the studies in the held-out test set. As shown in Supplementary Figure 5, Model-C+R improves patient stratification across PD-L1 expression cutoffs.

#### 3.4 SHAP Analysis

SHAP provides local interpretability via Shapley values which enable quantification of the contribution of each feature to the prediction problem. SHAP summary plots depicts one row for a covariate and a reference risk vertical line. Dots to the left of that line represent patients with lower risk than the reference while those on the right represent patients with higher risk.

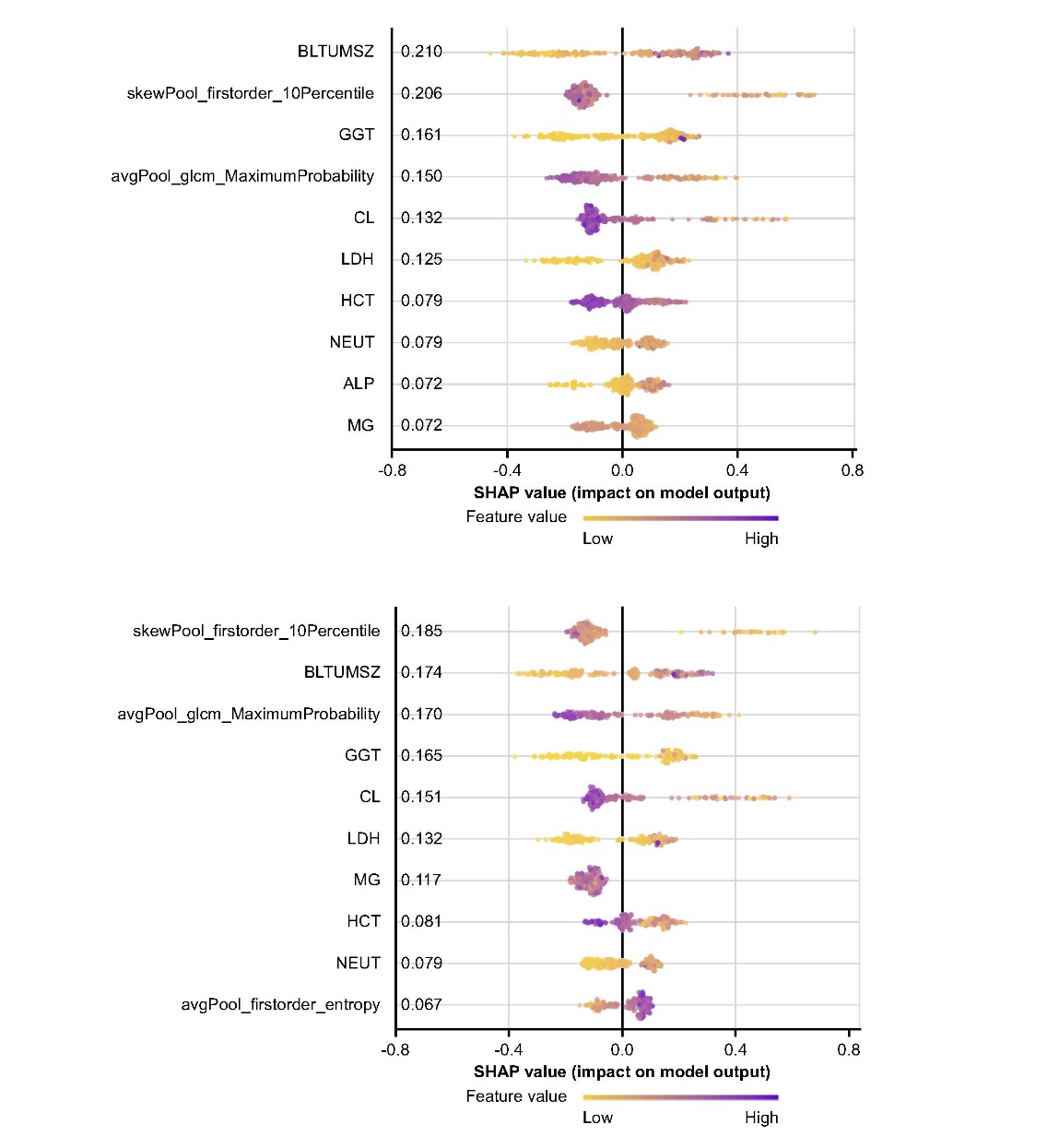

Supplementary Figure 6. SHAP summary plot highlighting the top-10 features used for the prediction in the test cohort (Top) and in the Independent study cohort (Bottom) using the clinical + radiomic model. ALP, alkaline phosphatase; BLTUMSZ, baseline tumor size; CL, chlorine; GGT, gamma glutamyl transferase; HCT, hematocrit; LDH, lactate dehydrogenase; MG, magnesium; NEUT, neutrophils; TSH, thyroid stimulating hormone.

Colors depict the feature value and the number next to a covariate indicates the mean SHAP values for that covariate. SHAP values are logarithms of hazard ratios and provide estimates of the size of effects. SHAP summary plots corresponding to the top-10 contributing features for each cohort were generated for Model-C+R in both the test cohort and independent study cohort and is shown in Supplementary Figure 6Error! No bookmark name given..

Supplementary Figure 7. SHAP response plot highlighting the top-6 features used for the prediction in the test cohort (Top) and in the Independent study cohort (Bottom) using the clinical + radiomic model. ALP, alkaline phosphatase; BLTUMSZ, baseline tumor size; CL, chlorine; GGT, gamma glutamyl transferase; HCT, hematocrit; LDH, lactate dehydrogenase; MG, magnesium; NEUT, neutrophils; TSH, thyroid stimulating hormone.

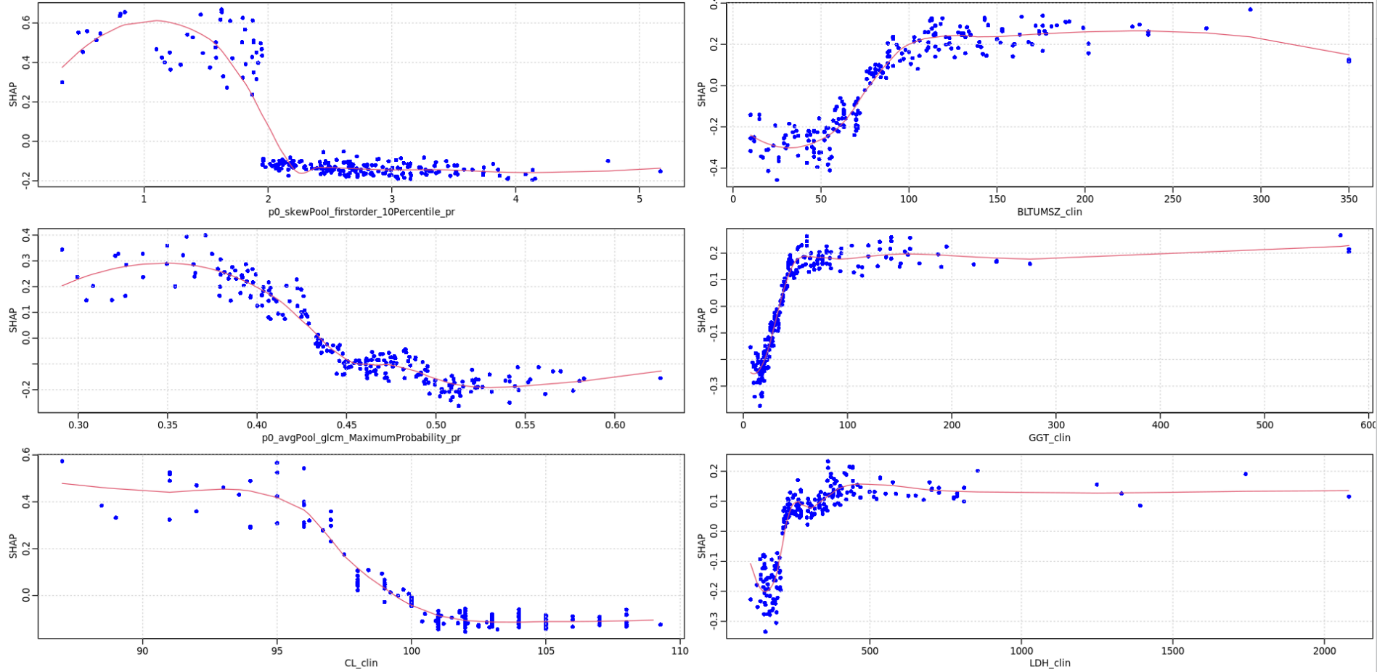

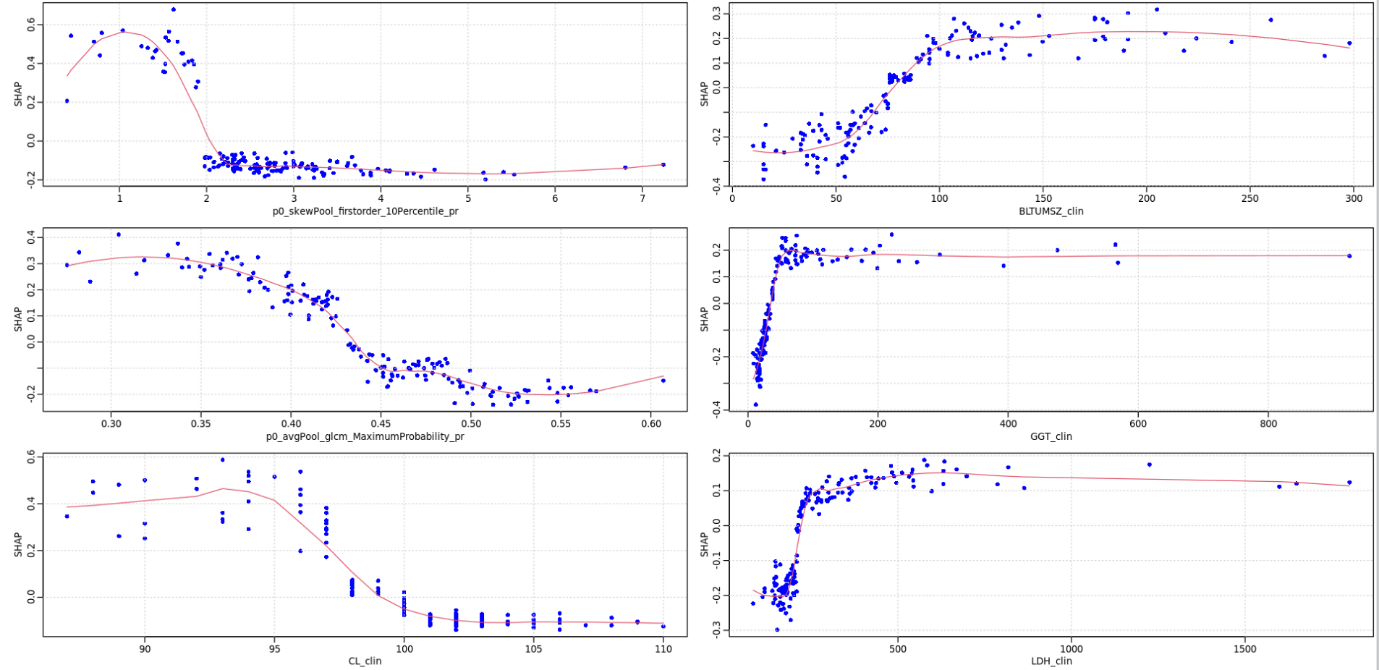

**a)**

**b)**

The summary statistic of the skewness and averaging of the first order and second order radiomic features respectively contributed to the model predictions. The 10^th^ percentile suggests that higher intensities in the lower 10% of voxel intensities would have a lower risk of survival. Similarly, higher number of occurrences of the most predominant pair of neighboring intensity values (gray-level cooccurrence matrix-maximum probability) suggests larger regions with similar intensities and a better survival on the treatment. From the comparison of these two plots, we can appreciate that the ranking of predictors is reasonably consistent and that top 6 predictors have mean SHAP values above 0.1 in both cases. Further interpretation of their effects was performed using SHAP response plots. The SHAP The reference risk level is indicated as “0.0” in the vertical axis. Patients at higher risk than the reference level are represented as points with positive SHAP values while those at lower risk are represented by points with negative risk level. A red spline curve is included in each plot to indicate the trend. Supplementary Figure 7 shows consistent trends in both cohorts. Further, a non-linear relationship was observed. The two top imaging features as well as baseline tumor size (BLTUMSZ) and chloride (CL) may be represented as dichotomous variables, with consistent thresholds across both cohorts. On the other hand, both gamma-glutamyl transferase (GGT) and lactate dehydrogenase (LDH) exhibit linear and saturated regimes.

#### 3.5 Model Reduction

Supplementary Figure 8. Correlation plots of the top features in a) test cohort and b) independent study cohort show that the imaging and clinical features are uncorrelated.

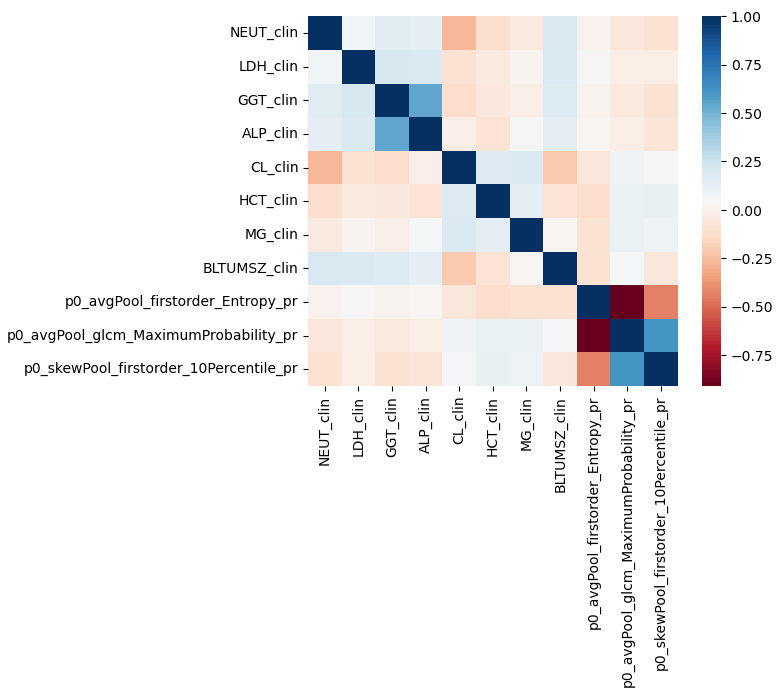

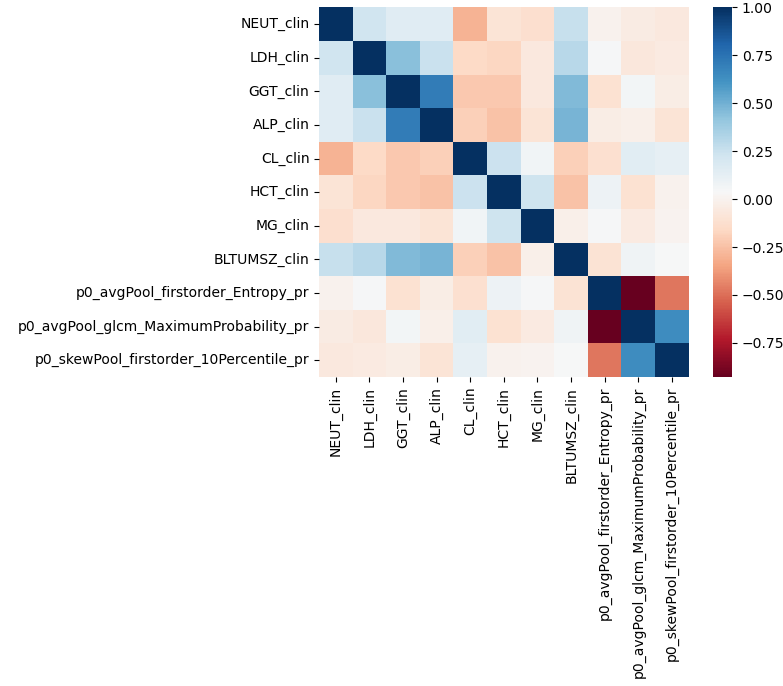

**a)**

**b)**

Prior to building a linear model, we observed that the imaging and clinical features were uncorrelated (shown in Supplementary Figure 8) supporting the build of a combined model. The top-6 predictors of overall survival were used to build a Cox model, which assumes linear relationships between the predictors and the response variable.

Supplementary Table 7. Significance of each variable in our first Cox model is shown. Please notice that the coefficients of these covariates represent increases per unit of the covariate, since they are continuous covariates. More clinically meaningful metrics will be obtained in Supplementary Table 9 & Supplementary Table 10, once we dichotomized some of these covariates.

|  | coef | exp(-coef) | se(coef) | z | Pr(>\|z\|) |
| --- | --- | --- | --- | --- | --- |
| skewPool_firstorder_10Percentile | -5.74E-02 | 9.44E-01 | 1.35E-01 | -0.425 | 0.671 |
| BLTUMSZ | 8.91E-03 | 1.01E+00 | 1.52E-03 | 5.852 | 4.85E-09*** |
| avgPool_glcm_MaximumProbability | -2.37E+00 | 9.35E-02 | 1.65E+00 | -1.434 | 0.152 |
| GGT | 1.08E-03 | 1.00E+00 | 8.28E-04 | 1.307 | 0.191 |
| CL | -1.45E-02 | 9.86E-01 | 2.26E-02 | -0.643 | 0.52 |
| LDH | -2.14E-05 | 1.00E+00 | 3.44E-04 | -0.062 | 0.95 |

Only baseline tumor size (BLTUMSZ) is considered to have a statistically significant linear relationship with overall survival (shown in Supplementary Table 7 & Supplementary Table 8). Without the SHAP response plots we may be tempted to believe that this implies that only BLTUMSZ is a relevant predictor of response, which we know not to be the case based on our SHAP analysis.

Supplementary Table 8. The confidence interval of each individual’s feature’s coefficient is shown below. The model has Concordance= 0.645 (se = 0.023 ), Likelihood ratio test= 39.21 on 6 df, p=6e-07, Wald test = 45.36 on 6 df, p=4e-08, Score (logrank) test = 46.06 on 6 df, p=3e-08. Please notice that the coefficients of these covariates represent increases per unit of the covariate, since they are continuous variables. More clinically meaningful metrics will be obtained in Supplementary Table 9 & Supplementary Table 10, once we dichotomized some of these covariates.

|  | coef | exp(-coef) | lower 0.95 | upper 0.95 |
| --- | --- | --- | --- | --- |
| skewPool_firstorder_10Percentile | 0.94418 | 1.0591 | 0.724604 | 1.23 |
| BLTUMSZ | 1.00895 | 0.9911 | 1.005941 | 1.012 |
| avgPool_glcm_MaximumProbability | 0.09354 | 10.6908 | 0.003666 | 2.386 |
| GGT | 1.00108 | 0.9989 | 0.999459 | 1.003 |
| CL | 0.98557 | 1.0146 | 0.942839 | 1.03 |
| LDH | 0.99998 | 1 | 0.999305 | 1.001 |

To represent properly the non-linear relationships observed in the SHAP response plots, we need to introduce appropriate transformations for the top 6 predictors:

- We built 4 new variables tum, skew, avg and cl as dichotomized versions of BLTUMSZ, skewPool_firstorder_10Percentile, avgPool_glcm_MaximumProbability and CL, respectively. For that purpose, we used 75, 2, 0.425 and 97.5 as thresholds, identified from the SHAP response plots.
- To represent the nonlinear relationships between GGT and LDH with overall survival, we used natural cubic splines with 3 degrees of freedoms.
- Further exploration (not shown here) using SHAP interaction plots suggested a possible interaction between skewPool_firstorder_10Percentile and GGT as well as between avg and LDH.

We are now in the position to build a new Cox model, which is still linear in its covariates, but which consider the non-linear relationships that exist between our top 6 predictors and the response variable as well as any relevant interaction that we have observed between predictors. This is an example of “model reduction,” which starts with a large and complex machine learning model and ends with a simple Cox model, which summarizes all the relevant information present in the datasets and it is easier to communicate to clinicians.

Supplementary Table 9. Significance of each variable in the Cox model with interactions between skew and GGT as well as avg and LDH is shown. For these dichotomized covariates, we cannot talk about hazard ratios (HR), which are more clinically meaningful metrics.

|  | coef | HR | se(coef) | z | Pr(>\|z\|) |
| --- | --- | --- | --- | --- | --- |
| tum | 4.74E-01 | 1.61E+00 | 1.78E-01 | 2.662 | 0.00777 ** |
| skew | 1.25E+00 | 3.50E+00 | 9.13E-01 | 1.373 | 0.16984 |
| ns(GGT, df = 3)1 | -9.87E-01 | 3.73E-01 | 1.84E+00 | -0.537 | 0.5914 |
| ns(GGT, df = 3)2 | 1.07E+01 | 4.28E+04 | 3.71E+00 | 2.876 | 0.00403 ** |
| ns(GGT, df = 3)3 | 1.65E+01 | 1.39E+07 | 6.60E+00 | 2.491 | 0.01273 * |
| cl | -4.40E-01 | 6.44E-01 | 2.29E-01 | -1.917 | 0.05527 . |
| avg | -2.16E+00 | 1.15E-01 | 7.89E-01 | -2.742 | 0.00611 ** |
| ns(LDH, df = 3)1 | -3.08E-01 | 7.35E-01 | 1.29E+00 | -0.239 | 0.81077 |
| ns(LDH, df = 3)2 | -4.03E-01 | 6.68E-01 | 2.10E+00 | -0.192 | 0.8479 |
| ns(LDH, df = 3)3 | 4.74E+00 | 1.15E+02 | 3.22E+00 | 1.474 | 0.14045 |
| skew:ns(GGT, df = 3)1 | 2.47E+00 | 1.19E+01 | 1.96E+00 | 1.259 | 0.20807 |
| skew:ns(GGT, df = 3)2 | -9.72E+00 | 6.01E-05 | 3.76E+00 | -2.585 | 0.00973 ** |
| skew:ns(GGT, df = 3)3 | -1.60E+01 | 1.13E-07 | 6.63E+00 | -2.413 | 0.01584 * |
| avg:ns(LDH, df = 3)1 | 1.22E+00 | 3.40E+00 | 1.46E+00 | 0.838 | 0.40227 |
| avg:ns(LDH, df = 3)2 | 1.02E+00 | 2.77E+00 | 2.34E+00 | 0.434 | 0.66409 |
| avg:ns(LDH, df = 3)3 | -5.33E+00 | 4.82E-03 | 3.42E+00 | -1.558 | 0.1193 |

Supplementary Table 9 & Supplementary Table 10 provide us with a reduced statistical representation from the main knowledge gathered by our analysis, that can be communicated more easily to clinicians. Our new model is statistically significant, with a concordance index of 0.64 and most of the top 6 predictors contribute in a statistically significant manner, except for CL which is nearly statistically significant with a p-value of 0.056. Dichotomized variables also allow us to compute hazard ratios (HRs) which provide more clinically meaningful metrics.

Finally, ANOVA tests, together with Akaike and Bayesian information criterion tests indicated that the two interactions were required to produce a better fit to the data.

Supplementary Table 10. The confidence interval of each individual’s feature’s hazard ratio is shown. Model Concordance= 0.644 (se = 0.023), Likelihood ratio test= 44.15 on 16 df, p=2e-04, Wald test = 52.63 on 16 df, p=9e-06, Score (logrank) test = 64.76 on 16 df, p=8e-08. For these dichotomized covariates, we cannot talk about hazard ratios (HR), which are more clinically meaningful metrics.

|  | HR | lower .95 | upper .95 |
| --- | --- | --- | --- |
| tum | 1.61E+00 | 1.13E+00 | 2.28E+00 |
| skew | 3.50E+00 | 5.85E-01 | 2.09E+01 |
| ns(GGT, df = 3)1 | 3.73E-01 | 1.02E-02 | 1.37E+01 |
| ns(GGT, df = 3)2 | 4.28E+04 | 2.99E+01 | 6.14E+07 |
| ns(GGT, df = 3)3 | 1.39E+07 | 3.34E+01 | 5.78E+12 |
| cl | 6.44E-01 | 4.11E-01 | 1.01E+00 |
| avg | 1.15E-01 | 2.45E-02 | 5.40E-01 |
| ns(LDH, df = 3)1 | 7.35E-01 | 5.90E-02 | 9.15E+00 |
| ns(LDH, df = 3)2 | 6.68E-01 | 1.08E-02 | 4.12E+01 |
| ns(LDH, df = 3)3 | 1.15E+02 | 2.10E-01 | 6.27E+04 |
| skew:ns(GGT, df = 3)1 | 1.19E+01 | 2.52E-01 | 5.57E+02 |
| skew:ns(GGT, df = 3)2 | 6.01E-05 | 3.79E-08 | 9.53E-02 |
| skew:ns(GGT, df = 3)3 | 1.13E-07 | 2.58E-13 | 4.98E-02 |
| avg:ns(LDH, df = 3)1 | 3.40E+00 | 1.94E-01 | 5.94E+01 |
| avg:ns(LDH, df = 3)2 | 2.77E+00 | 2.80E-02 | 2.74E+02 |
| avg:ns(LDH, df = 3)3 | 4.82E-03 | 5.87E-06 | 3.97E+00 |

### 4 References

1.	Shen L, Tao G, Fu H, Liu X, Ye X, Ye JJZZlzz. Predicting response to non-small cell lung cancer immunotherapy using pre-treatment contrast-enhanced CT texture-based classification. 2021;43(5):541-5.

2.	Wen Q, Yang Z, Dai H, Feng A, Li QJFio. Radiomics Study for Predicting the Expression of PD-L1 and Tumor Mutation Burden in Non-Small Cell Lung Cancer Based on CT Images and Clinicopathological Features. 2021:3082.

3.	Yang Y, Yang J, Shen L, Chen J, Xia L, Ni B, et al. A multi-omics-based serial deep learning approach to predict clinical outcomes of single-agent anti-PD-1/PD-L1 immunotherapy in advanced stage non-small-cell lung cancer. 2021;13(2):743.

4.	Sun Z, Hu S, Ge Y, Wang J, Duan S, Song J, et al. Radiomics study for predicting the expression of PD-L1 in non-small cell lung cancer based on CT images and clinicopathologic features. 2020;28(3):449-59.

5.	Vaidya P, Bera K, Patil PD, Gupta A, Jain P, Alilou M, et al. Novel, non-invasive imaging approach to identify patients with advanced non-small cell lung cancer at risk of hyperprogressive disease with immune checkpoint blockade. 2020;8(2).

6.	Liu Y, Wu M, Zhang Y, Luo Y, He S, Wang Y, et al. Imaging biomarkers to predict and evaluate the effectiveness of immunotherapy in advanced non-small-cell lung cancer. 2021;11:773.

7.	Yang B, Zhou L, Zhong J, Lv T, Li A, Ma L, et al. Combination of computed tomography imaging-based radiomics and clinicopathological characteristics for predicting the clinical benefits of immune checkpoint inhibitors in lung cancer. 2021;22(1):1-15.
